# Aperiodic activity as a central neural feature of hypnotic susceptibility outside of hypnosis

**DOI:** 10.1101/2023.11.16.567097

**Authors:** Mathieu Landry, Jason da Silva Castanheira, Catherine Boisvert, Floriane Rousseaux, Jérôme Sackur, Amir Raz, Philippe Richebé, David Ogez, Pierre Rainville, Karim Jerbi

## Abstract

How well a person responds to hypnosis is a stable trait, which exhibits considerable inter-individual diversity across the general population. Yet, its neural underpinning remains elusive. Here, we address this gap by combining EEG data, multivariate statistics, and machine learning in order to identify brain patterns that differentiate between individuals high and low in susceptibility to hypnosis. In particular, we computed the periodic and aperiodic components of the EEG power spectrum, as well as graph theoretical measures derived from functional connectivity, from data acquired at rest (pre-induction) and under hypnosis (post-induction). We found that the 1/f slope of the EEG spectrum at rest was the best predictor of hypnotic susceptibility. Our findings support the idea that hypnotic susceptibility is a trait linked to the balance of cortical excitation and inhibition at baseline and offers novel perspectives on the neural foundations of hypnotic susceptibility. Future work can explore the contribution of background 1/f activity as a novel target to distinguish the responsiveness of individuals to hypnosis at baseline in the clinic.

**Significance Statement:** Hypnotic phenomena reflect the ability to alter one’s subjective experiences based on targeted verbal suggestions. This ability varies greatly in the population. The brain correlates to explain this variability remain elusive. Addressing this gap, our study employs machine learning to predict hypnotic susceptibility. By recording electroencephalography (EEG) before and after a hypnotic induction and analyzing diverse neurophysiological features, we were able to determine that several features differentiate between high and low hypnotic susceptible individuals both at baseline and during hypnosis. Our analysis revealed that the paramount discriminative feature is non-oscillatory EEG activity before the induction—a new finding in the field. This outcome aligns with the idea that hypnotic susceptibility represents a latent trait observable through a plain five-minutes resting-state EEG.

## Introduction

Hypnosis involves using targeted verbal suggestions that engage top-down mental processes to modify perception, cognition, ideomotor actions, and emotional responses. (Terhune et al., 2017). In this way, hypnosis enables individuals to gain control over mental processes that are otherwise difficult to control (Lifshitz et al., 2013). Beyond the response to verbal suggestions, hypnotic phenomena are also characterized by spontaneous changes in phenomenology, including heightened feelings of relaxation and absorption, shifts of volitional control, as well as temporal distortions (Kumar & Pekala, 1989; Pekala, 2015; Rainville & Price, 2003; Timmermann et al., 2023; Weitzenhoffer, 1980). Accordingly, hypnotic phenomena are multifaceted and hardly reducible to a single dimension or feature.

The neuroscientific landscape of hypnosis reflects this intricate complexity, with research spanning various hypotheses, methodologies, and frameworks (Jamieson, 2007). This diversity makes it challenging to identify commonalities across studies (De Pascalis, 2023; Landry et al., 2017; Vanhaudenhuyse et al., 2014). In the context of electrophysiological recordings, previous work reveals that several aspects of the neural signal relate to hypnotic phenomena, including the spectral power (e.g., De Pascalis & Penna, 1990), evoked responses (e.g., Spiegel et al., 1985), its fractal component (Lee et al., 2007), the complexity of broadband signals (Rho et al., 2021), shifts in connectivity patterns (e.g., Jamieson & Burgess, 2014), as well as changes in the intricate organization of network topology (Panda et al., 2023). Results based on the spectral features include findings in the alpha-band and theta-band activity, whereas connectivity patterns have been found across several bands (De Pascalis, 2024b). Neuroimaging studies also highlights the involvement of various brain regions, including modulations of resting-state networks—such as the central executive, the salience, and the default networks (Landry et al., 2017). This variety of findings in the literature reflects the inherent complexity of hypnosis and demonstrates its multifaceted nature. Taken together, the existing body of work not only highlights the diverse methodologies and perspectives employed to probe hypnosis but also points to the intricate relationship between hypnotic phenomena and various underlying cognitive and neural mechanisms (Landry & Rainville, 2024).

To tackle this complexity, neuroscientific research largely centers on the exploration of three key variables: Susceptibility to hypnotic suggestions, the induction procedure, and the impact of suggestions (Landry & Raz, 2015). Susceptibility to hypnotic suggestions refers to a stable latent trait that primarily signifies the ability to effectively respond to direct verbal suggestions within controlled and standardized settings (Laurence et al., 2008). This entails that hypnotic responding represents at least in part an aptitude, much like athletic performances reflect a physical aptitude to perform at certain level. Notably, susceptibility to hypnotic suggestions serves as a predictive indicator across both experimental and clinical contexts of hypnosis (Barnier et al., 2014). In turn, the induction procedure constitutes an integral part of most hypnotic protocols, designed to enhance an individual’s readiness for verbal suggestions through a trance-like mental state (Yapko, 2015). This procedure primarily involves guiding individuals towards deep absorption and relaxation with the aim of facilitating the hypnotic response.

Markedly, the induction procedure is related with spontaneous changes in phenomenology (Kumar & Pekala, 1989). However, it remains uncertain how these modifications ultimately contribute to the hypnotic response (Terhune & Cardeña, 2016).

The primary objective of this study is to elucidate the neural dynamics underlying susceptibility to hypnosis, both before and after hypnotic induction. Building on prior research, our study examines spectral features, including aperiodic activity—a novel feature in the neurophysiological investigation of hypnosis. The aperiodic component of the neural power spectrum corresponds to scale-free background activity that approximates a power-law distribution (*P*∝1/*f^β^*), whereas the periodic component denotes discernible peaks in the frequency domain (Donoghue et al., 2022; Wilson et al., 2022). Both the aperiodic and periodic elements of the power spectrum hold relevance across a wide range of psychological phenomena, spanning sensations and cognition, developmental trajectories and aging, as well as clinical and pharmacologically-induced mental states, and altered states of consciousness (Gerster et al., 2022; Gyurkovics et al., 2022; He, 2014; Lopes da Silva, 2013; Maschke et al., 2023; Pei et al., 2023). Our approach also extends to analyzing network topology using graph theoretical approach, further aligning with established findings in the field (Panda et al., 2023).

Previous research proposes that the capacity for hypnotic response may reflect a latent trait rooted in neural processes, implying that hypnotic susceptibility corresponds to intrinsic neural traits (De Pascalis, 2024a). Here, we used an experimental approach that includes 5 minutes resting-state electroencephalography (EEG) both before and after the hypnotic induction to evaluate this perspective and determine whether the neural basis of hypnotic susceptibility is more pronounced inside or outside of hypnosis Observing the primary neural feature outside of hypnosis would support the concept of a latent neural trait. We accomplish this through the integration of high-density EEG, advanced multivariate pattern classification techniques, and machine learning methodologies. Compared to univariate approaches, multivariate approaches increase sensitivity in detecting effects based on distributed patterns across multidimensional spaces (Haynes & Rees, 2006). We accordingly aimed to assess which neural features could be used to discriminate between individuals with high (HHSIs) and low hypnotic susceptibility (LHSIs) both at baseline--prior to the hypnotic induction--and based on the hypnotic effect (post-minus pre-induction: Δinduction; **Figure 1**).

**Figure 1.**
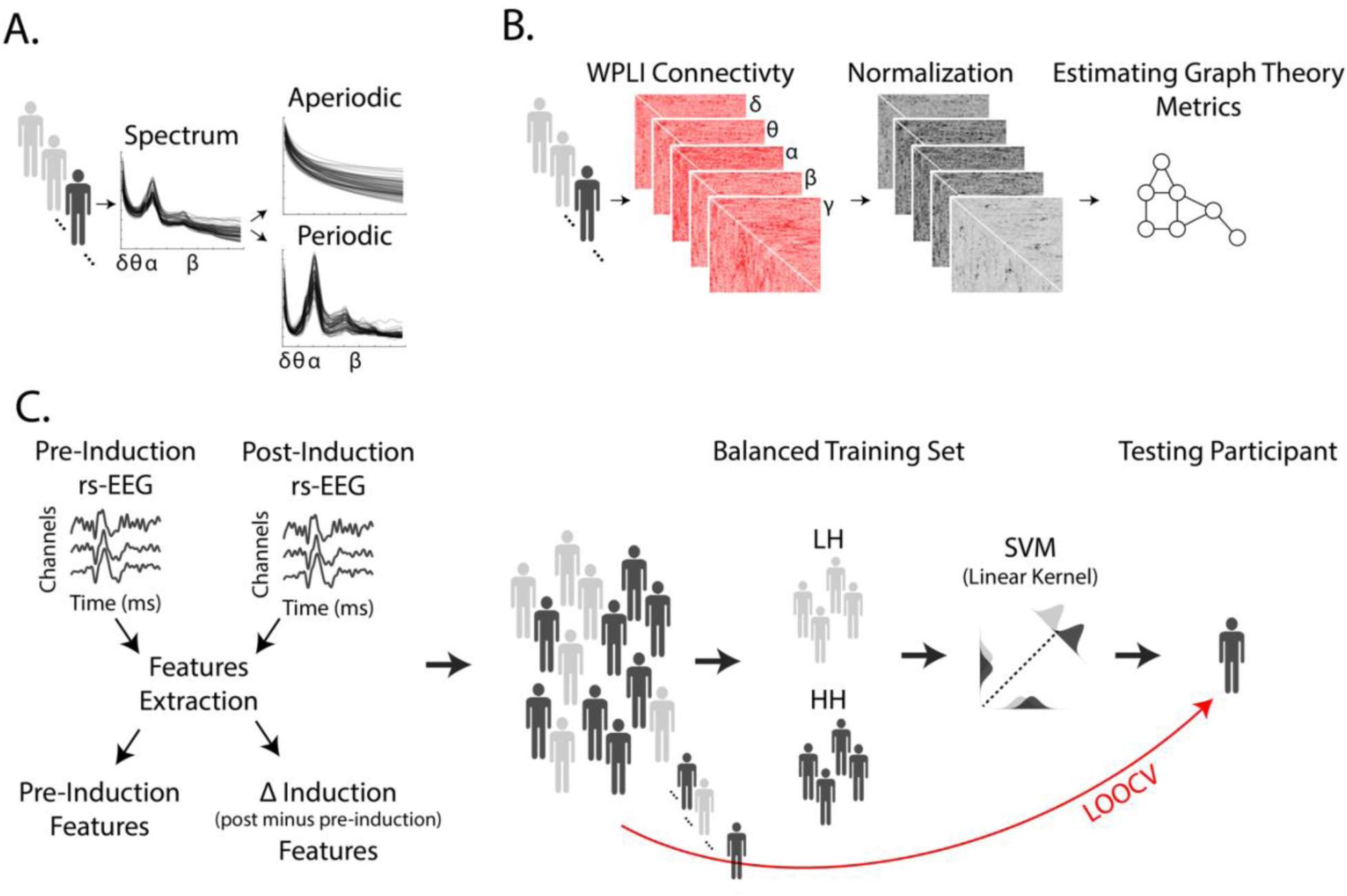
Analysis pipeline for extracting power spectrum and network topology features and performing multivariate pattern analysis with leave-one-out cross-validation. **A.** Aperiodic and periodic components of the power spectrum were estimated for each participant using 5 minutes of eye-closed rs-EEG data, both pre- and Δinduction. **B.** Connectivity patterns between EEG channels were calculated using weighted Phase Lag Index (wPLI) across delta, theta, alpha, beta, and gamma frequencies. The connectivity matrices were normalized from 0 to 1 (Rubinov & Sporns, 2011). Clustering coefficient, global efficiency, and modularity were computed using graph theory for each network. **C.** High and low hypnotic susceptible individuals were classified based on these metrics using multivariate pattern analysis with leave-one-out cross-validation (LOOCV) employing linear Support Vector Machine (SVM). EEG channels were included as features of this linear model. Each training iteration included balanced classes of low (light grey) and high (dark grey) hypnotic susceptible individuals, while one participant was left out for validation.

The present study addresses two key research questions. Firstly, we aim to identify the neural characteristics associated with hypnotic susceptibility before and after hypnotic induction. Given the complexity of hypnotic phenomena and previous research findings, we hypothesize that various attributes captured by EEG will be directly relevant to assessing hypnotic susceptibility, both prior to and following the induction procedure. Secondly, we seek to determine the top neural feature for discriminating high versus low hypnotic susceptible individuals and then examine whether this feature is observable inside or outside hypnosis. The presence of this principal neural feature outside of hypnosis would support the idea that susceptibility to hypnosis is a stable latent trait, which can be assessed independently of hypnotic induction.

## Methods

### Participants

Our recruitment procedure comprised two stages. First, potential participants were invited to a pre-screening session, during which we assessed their hypnotic susceptibility using the Harvard Group Scale of Hypnotic Susceptibility, Form A (HGSHS:A; Shor & Orne, 1962). Subsequently, we selected 75 individuals from this initial pool to undergo EEG recordings. For the present study, we focused on neural activity data from 40 individuals (27 females, mean age of 21.98 y.o. (st.dev. = 4.41)) based on their HGSHS:A scores. Specifically, we included individuals who scored above 8, indicating high hypnotic susceptibility (HHSIs), and individuals who scored below 4, indicating low hypnotic susceptibility (LHSIs). This work aligns with previous studies focusing on high and low hypnotic susceptible individuals to maximize the detection of the effect. We Participants were not informed of their scores. Participants who completed the second part of our recruitment procedure received monetary compensation. All study procedures were approved by the local institutional review board. Participants provided written consent.

The sample size for our study was established based on prior research that explored the impact of hypnotic susceptibility on spectral features and connectivity measures in EEG recordings, with these studies reporting statistically significant findings. Specifically, research has indicated significant influences of hypnotic susceptibility within the alpha-band and theta-band during resting-state EEG activity. For instance, Palumbo De Pascalis and Palumbo (1986) included 20 HHSIs and 20 LHSIs. Graffin et al. (1995) worked with 14 HHSIs and 13 LHSIs, while Williams and Gruzelier (2001) studied 8 HHSIs and 8 LHSIs. Crawford et al. (1996) examined 15 HHSIs and 16 LHSIs. Furthermore, De Pascalis and Penna (1990) with recordings done with 19 HHSIs and 20 LHSIs and De Pascalis (1993) with 9 HHSIs and 10 LHSIs report in the gamma range. However, it is important to note that Jamieson and Burgess (2014) did not observe a significant effect of hypnotic susceptibility across different frequency bands with their sample of 12 HHSIs and 11 LHSIs. For connectivity patterns,

### Procedure

The assessment of hypnotic susceptibility was conducted following the standard procedure of the HGSHS:A in a separate experimental session on a different day. The experimental procedure consisted of a 5-minute EEG that was recorded at baseline (i.e., pre-induction period), followed by the hypnotic induction procedure, and subsequently, another 5-minute EEG recording.

During the EEG recording, participants were seated in a quiet and dimly lit room. The experimenter, who conducted the induction procedure, positioned themselves approximately one meter behind the participants throughout the experiment. We recorded the baseline EEG for all participants before the induction-related EEG, instead of counterbalancing our experimental conditions. This approach prevented contamination (i.e., carry-over effects) of the baseline period by the hypnotic induction, as it is challenging to determine the duration required for the effects of hypnosis to wash out after the de-induction. Additionally, since the objective of the current study was to differentiate highly hypnotic susceptible individuals from those with low susceptibility, the order of the baseline and induction-related EEG recordings was deemed irrelevant. In sum, for each participant, we first recorded 5 minutes EEG resting-state activity outside of hypnosis at baseline, and then again 5 minutes EEG resting-state activity following the hypnotic induction procedure while participants were in hypnosis.

Participants were instructed by the experimenter to close their eyes while remaining vigilant during the baseline recording. The recording lasted 5 minutes. Immediately after, the experimenter indicated to participants that the induction procedure would begin, verified that participants were ready, and then read a script for the induction (see supplementary material). The induction procedure included instructions for participants to close their eyes, and the 5-minute EEG recording was conducted after the completion of the induction script. Three different experimenters trained in hypnosis contributed to data collection and performed the induction. We did not collect subjective reports concerning the success of the induction. Following previous work, we assumed that the induction procedure would differentially impact LHSIs and HHSIs (for review, see Landry et al., 2017). Our results corroborate this assumption.

### Electroencephalography

We recorded EEG signals using 64 Ag/AgCl active electrodes at a sampling rate of 1000 Hz (ActiCap System; Brain Products GmbH; Gilching, Germany). We positioned additional bipolar electrodes at the outer canthi, as well as the superior and inferior orbits of the left eye. We kept impedances of all electrodes below 10 kΩ, while all electrophysiological signals were amplified (ActiChamp System; BrainPProducts GmbH; Gilching, Germany). Electrode impedences were kept below 10 kΩ. Data was collected online at 1000Hz and down sampled to 250Hz offline for data analysis. Data acquisition was done via Brain Recorder (Brain Products Inc., GmbH) while pre-processing and data analysis were completed through Brain Vision Analyzer, Brainstorm (Tadel et al., 2011), and custom MATLAB scripts (version R2022A; MathWorks Inc, MA). Bad electrodes were topographically interpolated using spherical splines (< 3% electrodes). EEG recordings were re-referenced to the average of all EEG electrodes offline. All EEG data were inspected for muscle-related, eye movement or technical artifacts, which were removed before continuing the analysis.

### Spectral features

Spectral features were computed individually for each channel using the *specparam* algorithm for spectral parametrization as implemented in Brainstorm (Figure 1A; Donoghue et al., 2020; Tadel et al., 2011). The *specparam* algorithm decomposes power spectra into periodic and aperiodic (1/f-like) components by fitting the power spectral density in log-log space. The periodic component is characterized by the center frequency, bandwidth, and relative amplitude of the oscillatory peak, while the aperiodic component is defined by the offset and slope of the power spectrum density. In our analysis, we employed the model-selection implementation of specparam to fit the power spectral density from 1 to 40Hz, considering up to 6 peaks, with a minimum peak height of 1db, a peak threshold of 2, a proximity threshold of 0.75, peak width limits between 0.5 to 12 Hz, and a fixed aperiodic mode. Goodness-of-fit was evaluated based on mean-square error, and no significant differences were observed between high and low hypnotic susceptible individuals for the pre-induction period and Δ induction. Specifically, AUC for classification was .51 (p > .5) for pre-induction EEG and .49 (for Δinduction on EEG using the multivariate pattern classification described below. We then extracted the power spectrum values for the periodic component of each EEG channel by subtracting the aperiodic component from the power spectrum density. Finally, we calculated the average spectrum values separately for delta (1 to 3.5Hz), theta (4 to 7.5Hz), alpha (8 to 13.5Hz), and beta (14 to 39Hz) broadband frequencies. In summary, our analysis encompassed six spectral features: the offset and slope of the aperiodic component, as well as corrected power for delta, theta, alpha, and beta oscillations.

### Features of network topology

To extract network topology features, we utilized graph theory metrics on sensor-level connectivity patterns, employing the Brain Connectivity Toolbox (Figure 1B; Rubinov & Sporns, 2010). Initially, we applied the Hilbert transformation to filter the EEG data at specific frequency bands, namely delta (1 to 3.5Hz), theta (4 to 7.5Hz), alpha (8 to 13.5Hz), beta (14 to 39Hz), and gamma (30 to 50Hz) frequencies. Subsequently, we estimated functional connectivity between all electrode pairs using wPLI to account for volume conduction effects on brain connectivity estimation (Vinck et al., 2011). Then, we opted to normalize the edges into weights ranging from 0 to 1 rather than converting the 64 by 64 functional network into sparse and binarized patterns (Rubinov & Sporns, 2011). This approach ensured the preservation of valuable information without loss due to subjective thresholding choices. From these weighted undirected connectivity matrices, we extracted three distinct graph measures for all frequency bands: The clustering coefficient, derived from the geometric mean of local triangle configurations associated with each node, which provides insight into the extent of node clustering (Onnela et al., 2005); global efficiency, measured as the inverse of the shortest path length, that quantifies the level of shared information across the entire network (Latora & Marchiori, 2001); and global modularity to estimate the degree to which networks are divided into communities (Newman, 2006). These graph theory metrics allowed us to evaluate network integration and segregation as a function of hypnotic induction and susceptibility to hypnotic suggestions. Our graph theoretical approach therefore yielded three separate neural features, clustering coefficient, global efficiency and global modularity across all broad frequency bands.

### Multivariate pattern analysis and leave-one-out cross-validation

We employed a multivariate pattern analysis combined with a leave-one-out cross-validation strategy to train and test linear support vector machine (SVM) models. The objective was to discriminate between high and low hypnotic susceptible individuals based on spectral and network topological features (**Figure 1C**). To minimize classification biases, we carefully balanced the classes in our training set, including 16 high and 16 low hypnotic susceptible individuals in each iteration (Thölke et al., 2022). In each iteration of the cross-validation process, one participant was left out for testing the SVM model, while we randomly selected 16 participants from each class for the training set. This procedure was repeated to ensure that each participant served as a test case 10 times for comprehensive cross-validation. The SVM linear model was trained using the 64 EEG channels as features for classification. We assessed the classification performance using the area under the curve (AUC).

The observed classification performance was compared to a surrogate distribution generated from 1000 random permutations. In each permutation, the class labels were randomly shuffled during the training process (Combrisson & Jerbi, 2015). Statistical significance was set at alpha = .05. The observed classification performance was considered statistically significant only if it outperformed 95% the performance of the upper-part (i.e., one-tail) surrogate distribution.

To facilitate interpretation, we further transformed the weights of the SVM classifiers based on the covariance of the training dataset (Haufe et al., 2014). These transformed weights were then averaged across all training iterations. We also extracted PSD and graph theoretical values for the channels where we observed the top 5% SVM weights to better understand how hypnotic susceptibility and the induction procedure impacted EEG.

We evaluated spectral features and clustering coefficients for pre-induction and Δinduction separately.

In addition to EEG, we applied our multivariate classification approach to vertical and horizontal electro-oculogram signals (VEOG and HEOG) to determine whether we could discriminate between HHSIs and LHSIs across both pre-induction and Δinduction separately. This classification procedure showed that we could not discriminate between HHSIs and LHSIs based on signals from eye movements.

### Logistic regression

We used logistic regression to determine whether graph theoretical metrics of global efficiency and modularity across delta, theta, alpha, beta and gamma bands activity to predict high hypnotic susceptibility. We opted for logistic regression over multivariate pattern analysis because these metrics are computed across all channels, rather than at the individual channel level. We evaluated two separate logistic regression models: A first one for the EEG recorded during the pre-induction period and second one for Δinduction. Both models were the following:

Logit(P(high hypnotic susceptibility)) = β0 + β1[Global Efficiency Delta band] + β2[Global Efficiency Theta band] + β4[Global Efficiency Alpha band] + β5[Global Efficiency Beta band] + β6[Global Efficiency Gamma band] + β7[Modularity Delta band] + β8[Modularity Theta band] + β9[Modularity Alpha band] + β10[Modularity Beta band] + β11[Modularity Gamma band]

In this way, we were able to identify individual neural features that discriminate between HHSIs and LHSIs.

### Feature comparisons

The previous analyses allowed us to identify spectral and network features from the pre-induction and Δinduction EEG that differentiate between high and low hypnotic susceptible individuals on their own. Next, we aimed to ascertain the top discriminant feature when we combine them. Our approach was twofold.

First, we used model comparisons. Using the previously described multivariate pattern classification analysis and leave-one-out cross-validation approach, we trained and tested SVM models that incorporated all possible combinations of features we previously identified as statistically significant on their own. This assessment was performed three times: once for the pre-induction EEG period, then for Δinduction EEG data, and thirdly for all possible combinations of significant features from both the pre-induction EEG and Δinduction. For each model, we calculated the AUC as a measure of classification performance. To compare the classification performance of all models, we utilized McNemar’s test (McNemar, 1947), which is a reliable method for comparing classification performances based on contingency tables (Dietterich, 1998). To account for multiple comparisons, we corrected the p-values from these tests using the false discovery rate correction method proposed by Benjamini and Hochberg (1995).

Second, we used feature ranking. Here included all significant features into SVM models and then ranked the absolute value of the SVM weights following the procedure from Haufe et al. (2014) described earlier. We applied this feature ranking procedure three times: once for all significant features from the pre-induction EEG, second for all significant features for Δinduction, and thirdly for significant features from both the pre-induction and Δinduction EEG.

## Results

We discriminated between and high and low hypnotic susceptible individuals using a linear support vector machine (SVM) with a leave-one-out cross validation procedure (**Figure 1C**). We first tested the classifier’s discrimination ability based on spectral features. These features included rhythmic activity across delta (1 to 3.5Hz), theta (4 to 7.5Hz), alpha (8 to 13.5Hz), and beta (14 to 39Hz) frequencies, as well as the aperiodic offset and exponent values from estimates of non-oscillatory activity (**Figure 1A**). See Methods for details.

### Spectral features

For the pre-induction period, our SVM classification yielded statistically significant results for the aperiodic exponent (**Figure 2A**; AUC = .69, p < .001). However, classification performance based on other spectral features, including oscillatory activity, did not reach significance (**Figure 2A**). We conducted further analysis to examine the weights of the classifiers, which provided insights into the specific contributions of the topographical features (**Figure 2A).** Notably, we observed a widespread neural pattern indicating that high hypnotic susceptible individuals exhibit a greater aperiodic exponent across the entire scalp. However, this pattern was particularly pronounced in the anterior part of the frontal site (**Figure 2A**; with the highest averaged coefficient value observed at channel AF8) and the right temporal region. We note that we did not observe any difference in aperiodic exponents between LHSIs and HHSIs for Δinduction (**Figure 1C**; **supplementary Figure 1C**). These findings highlight the importance of aperiodic activity relative to hypnotic susceptibility, while the induction procedure does not alter this neural feature.

**Figure 2.**
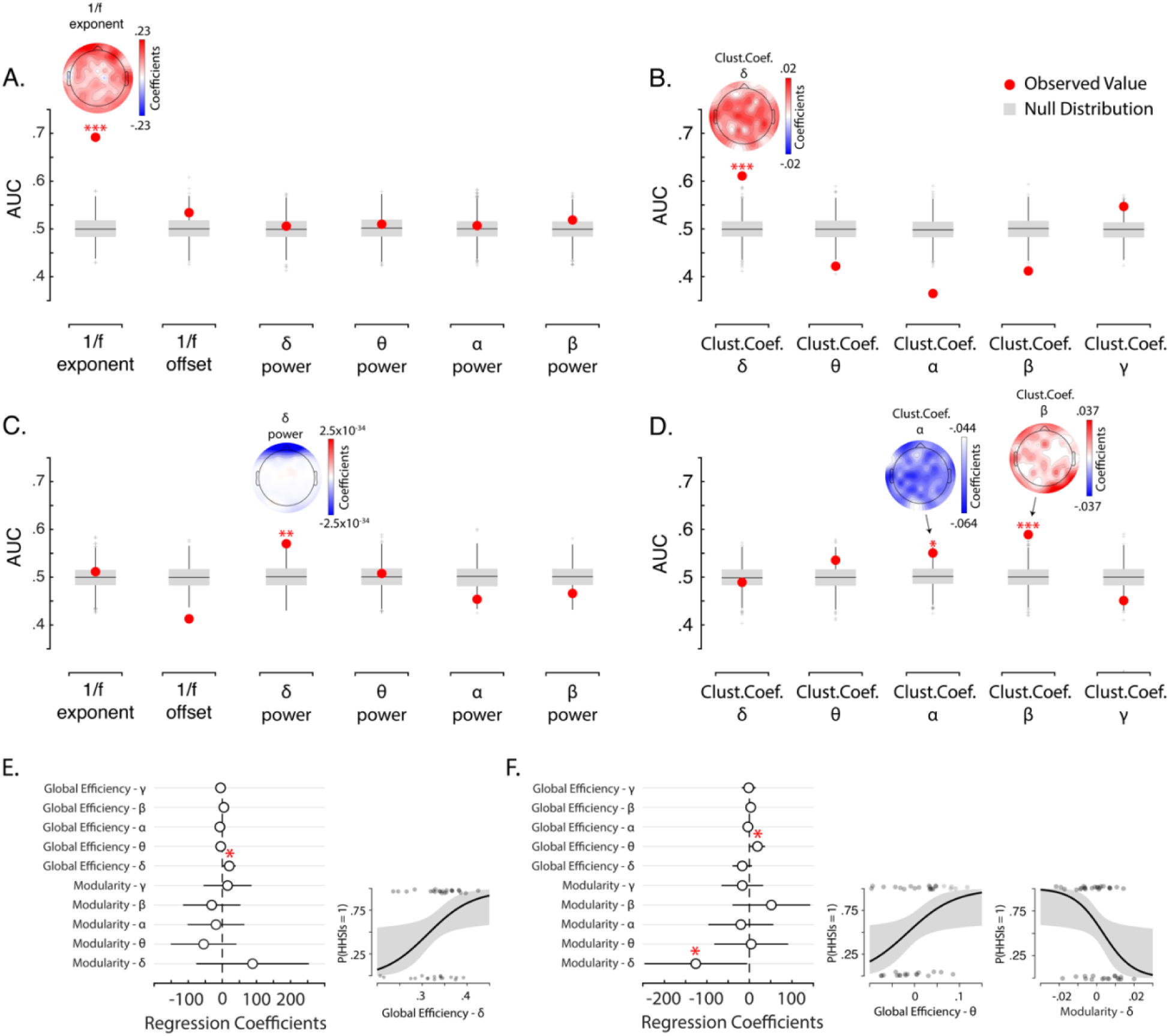
Panels A and B show analysis for pre-induction EEG, whereas panels C and D show analysis for Δinduction EEG. Panels A, B, C and D show AUC values for assessing classification performances of SVM linear models for accurately classifying high and low hypnotic susceptible individuals in LOOVC based on aperiodic and periodic features of the power spectrum during the EEG pre-induction period (A) and the Δinduction period (C), as well as graph theoretical clustering coefficients calculated from pairwise wPLI across frequency bands during the EEG pre-induction period (A) and for Δinduction (D). Red dots show observed AUC values. Boxplot shows surrogate null distributions based on random permutations where we shuffled the labels during training. The bottom and top whiskers indicate the first and third quartiles, respectively. Three asterisks indicate statistical significance at p < .001, two asterisks at p < .01, one asterisk at p < .05. Topographies show the averaged coefficient values of the SVM models from all iterations and LOOVC approach for the neural features that can discriminate high and low hypnotic susceptible individuals better than chance-level. Panel E shows the coefficient values and 95 C.I. from the logistic regression model assessing if global efficiency and modularity for pre-induction EEG across all frequency bands predict hypnotic susceptibility. Panel F show likelihood estimates for significant coefficients, global efficiency in the delta frequency band. Each dot represents individual data points and the grey area represents 95% C.I. The sigmoid function shows likelihood estimates for significant coefficients. Each dot represents individual data points, and the grey area represents 95% C.I. Red asterisks indicate significant coefficient at p < .05.

In contrast, periodic delta activity distinguished between LHSIs and HHSIs for Δinduction (**Figure 2C**; AUC = .57, p < .01). The corresponding topography of the transformed classifier’s weights highlights the frontal channels (**Figure 2C**). We note that during the post-induction, HHSIs and LHSIs exhibit similar delta power values, whereas HHSIs appear to show greater delta power value during pre-induction period (**supplementary Figure 1D-F**). Hence, our findings reveal that the induction has a greater impact on rhythmic delta band activity for HHSIs compared to LHSIs.

Note we assessed the contribution of ocular artifacts to participant classification by defining classifiers based on electro-oculogram signals. This analysis revealed we could not discriminate between HHSIs and LHSIs.

### Network topology

We then evaluated the classifier’s ability to discriminate between high and low hypnotic susceptible individuals based on graph theory clustering coefficients from channel-wise connectivity patterns as neural features (**Figure 1B**). Connectivity was estimated applying weighted Phase Lag Index (Vinck et al., 2011) based on neural activity across delta (1 to 3.5Hz), theta (4 to 7.5Hz), alpha (8 to 13.5Hz), beta (14 to 39Hz), and gamma (30 to 50Hz) frequencies.

We first examined whether classifiers could distinguish between LHSIs and HHSIs individuals based on clustering coefficients. In the pre-induction EEG recording, we found that the classification performance was statistically significant for clustering coefficient in the delta frequencies (**Figure 2B**; AUC value = .61, p < .001). Analyzing the weights of the classifiers, we observed widespread higher clustering coefficients in high hypnotic susceptible individuals, slightly concentrated in the central and bilateral temporal regions (**Figure 2B**). We observed higher coefficient values for HHSIs during the pre-induction period (**supplementary Figure 2A**). No other classification patterns reached statistical significance for the pre-induction EEG data (**Figure 2B**). These findings therefore provide further evidence of distinct neural characteristics associated with hypnotic susceptibility in the pre-induction state.

For Δinduction, we found that the classification performance was statistically significant in the alpha and beta frequency ranges (**Figure 2D**; AUC value for alpha-band activity = .55, p < .05; AUC value for beta-band activity = .59, p < .001). We observed opposite patterns between alpha-band and beta-band clustering coefficients, wherein HHSIs showed more pronounced decrease for the former and increase for the latter following the induction procedure (**Figure 2D**; **supplementary Figure 2**). These findings provide evidence that the induction process yields greater changes in clustering coefficients for individuals with high hypnotic susceptibility compared to those with low hypnotic susceptibility.

Next, we used logistic regression analyses to examine the effects of hypnotic susceptibility on global efficiency and modularity at the network level. Here we included both global efficiency and modularity estimates across broadband frequencies (delta, theta, alpha, beta, gamma) as predictors in our model for predicting hypnotic susceptibility. For the pre-induction EEG data, the logistic regression model (VIFs < 2.8) indicates that global efficiency in delta band activity was a statistically reliable predictor of hypnotic susceptibility (**Figure 2E**). Here, greater global efficiency predicts high hypnotic susceptibility during the pre-induction EEG period (β = 20.16, 95% C.I. [1.3, 39.02]; **Figure 2E**). For Δinduction, the logistic regression model (VIFs < 2.5) revealed that global efficiency in the theta frequencies (β = 18.9, 95% C.I. [.92, 36.99]) and modularity in the delta frequencies (β = -126.01, 95% C.I. [-246.67, -5.35]) were statistically significant predictors (**Figure 2F**). Specifically, the induction procedure was associated with increased global efficiency in the theta frequencies (**Figure 2F**) and decreased modularity (**Figure 2F**) in the delta frequencies in individuals with high hypnotic susceptibility compared to those with low hypnotic susceptibility.

### Model comparisons

We used model comparisons to identify the most dominating feature among those we previously identified as statistically significant. Our approach involved systematically evaluating all combinations of statistically significant spectral features and graph theory metrics and then assessing the classification performance of these models. We compared the performance of all models in a pairwise manner using the McNemar test and corrected the p-values for multiple comparisons using the false discovery rate (FDR) method. This approach allowed us to assess which combination of variables provided the best classification performance for each period.

Our results show that the models incorporating the aperiodic exponent consistently outperformed the other models, as indicated by higher AUC values for all these models (**Figure 3A**). The McNemar’s Chi-square statistics confirmed the advantage of these models over classification performance (**Supplementary Figure 3A**).

**Figure 3.**
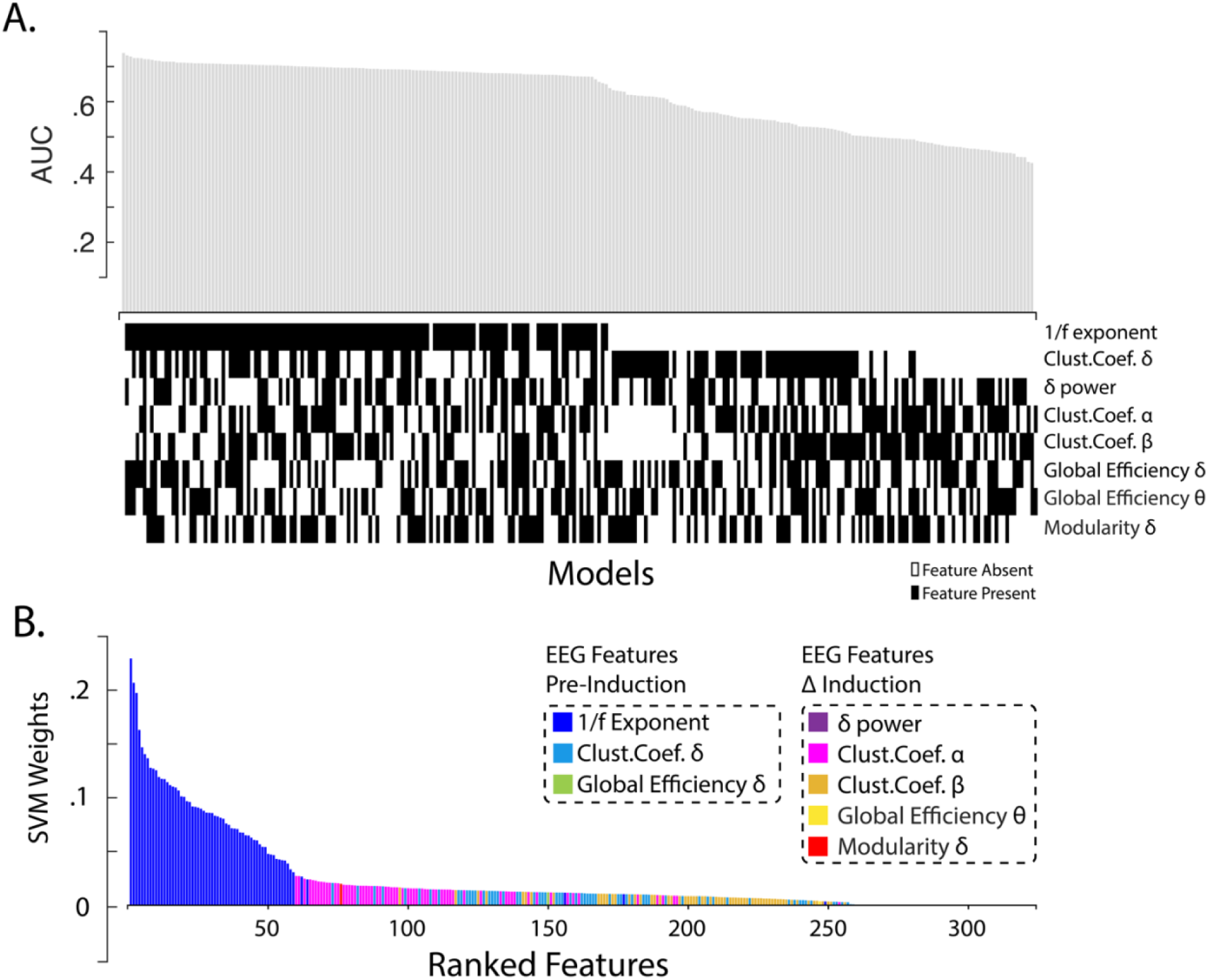
Panel A shows AUC values for each model that were evaluated for the discrimination of high versus low hypnotic susceptible individuals based on neural features from EEG from the pre-induction period and Δinduction. All features had been identified via previous analyses. The x-axis shows the neural features that were included in each model wherein a black square indicates that the neural feature was included in the model, and a white square indicates that the feature was not included in the model. Weights of SVM classifiers trained to differentiate high and low hypnotic susceptibility using EEG features from both pre-induction EEG period and Δinduction. Features were identified in preceding analyses. The x-axis ranks neural features in descending order, while the y-axis displays the absolute values of the SVM weights.

### Neural feature ranking

We sought to rank the pertinent neural features by their SVM weights based on the SVM model incorporating all relevant features. Examining feature ranking for all significant features from both the pre-induction period and Δinduction, the model performed above chance-level (AUC value = .69, p < .001). Again, the exponent of aperiodic component for the pre-induction resting-state period was the most discriminant feature, followed by the clustering coefficient of alpha-band activity in EEG pertaining to the effect of the induction procedure (**Figure 3B**). Collectively, these analyses highlight the centrality of aperiodic EEG activity for discriminating hypnotic susceptibility.

## Discussion

The current study provides a comprehensive investigation into the neural correlates of an individual’s receptiveness to hypnotic suggestion. Our findings confirmed our first hypothesis and show that the neural basis of hypnotic susceptibility is complex and multifaceted, as several neural features allowed us to discriminate between HHSIs and LHSIs. Regarding our second hypothesis, our results are consistent with the idea that hypnotic susceptibility reflects a latent trait observable at the neural level. Here, we found that the top neural feature for predicting individual’s hypnotic susceptibility is observable from their ongoing brain data recorded before hypnotic induction, principally from the aperiodic component of brain activity. These findings complement earlier observations from functional neuroimaging studies that relate neural functioning to hypnotic susceptibility (Hoeft et al., 2012; Huber et al., 2014), and provide an electrophysiological account for the intriguing observation that susceptibility to hypnotic suggestion is a predisposition that varies across the general population.

Our findings reveal that neural responses to hypnotic induction differ significantly between HHSIs and LHSIs across several frequency bands. Notably, we observed that rhythmic delta band activity was distinctly pronounced among individuals high on hypnotic susceptibility during hypnosis. This observation aligns with a recent study that reported similar findings in high hypnotic susceptible individuals prone to experiencing dissociation (Taghilou et al., In press). Still, it remains unclear how modulations of delta-band activity contribute to hypnotic phenomena. In contrast, we found no significant effects of hypnotic induction on theta and alpha power—two neural patterns frequently associated with hypnosis in prior studies (De Pascalis, 2024b). Changes in theta oscillations during hypnosis are thought to reflect mental processes that facilitate hypnotic responding via coupling with gamma band activity (Jensen et al., 2015). This coupling is pivotal for declarative memory, a cognitive process purportedly involved in hypnosis according to this viewpoint. Similarly, alpha-band activity is likely indicative of changes in relaxation levels during hypnosis, consistent with increasing evidence that links alpha oscillations to mental relaxation (Sugimoto et al., 2024). However, our findings challenge the prevailing notion that theta and alpha oscillations are central to hypnotic phenomena, as we observed no differential patterns in these neural components between high hypnotic susceptibility individuals (HHSIs) and low hypnotic susceptibility individuals (LHSIs). Consequently, the role of theta and alpha bands in the emergence of hypnosis remains uncertain.

Recent work suggests that hypnotic induction induces polyrhythmic changes in network topology (Panda et al., 2023). At the psychological level, the induction procedure is associated with increased mental absorption, feelings of relaxation, and preparation for hypnotic responding (Terhune & Cardeña, 2016). The effects we observed relative to modifications in network topologies are therefore likely indicative of the differential expression of these mental states across HHSIs and LHSIs. However, we did not collect phenomenological reports related to the induction procedure, which limits our ability to relate these neural dynamics to specific mental states. Interestingly, analogous shifts in network structures across delta, alpha, and beta frequencies have also been observed when comparing neural patterns of novice and expert meditators (Lardone et al., 2022; van Lutterveld et al., 2017). This observation resonates with the hypothesis suggesting a convergence between hypnosis and meditative practices (Raz & Lifshitz, 2016). Indeed, both practices encompass the involvement of attention resources and the recruitment of overlapping brain networks (Landry & Raz, 2016; Lifshitz et al., 2019).

Importantly, our results reveal that the aperiodic component of brain activity outside of hypnosis (i.e., at baseline) stands out as the primary neural feature that contributes most to the classification. This outcome supports the idea that hypnotic susceptibility reflects a cerebral predisposition and brings about the possibility of assessing it using EEG instead of using standardized protocols (Baghdadi & Nasrabadi, 2012). To some extent, this result challenges the assumption that the primary neural signature indicative of hypnotic susceptibility should become apparent following hypnotic induction. However, it remains important to note that participants were aware of the experimental context, potentially introducing expectation effects into our findings. Prior research emphasizes how expectations can significantly influence hypnotic responses (Gandhi & Oakley, 2005). Thus, while the prominent neural feature was observed outside of hypnosis, differing expectation levels between HHSIs and LHSIs during baseline EEG recordings may have influenced these results. Still, mounting evidence proposes that ability for hypnotic responding is best captured by interrelated components characterized by a structure comprising a core ability that superordinate secondary ones (Barnier et al., 2022). One may therefore conjecture that the centrality of the aperiodic slope for differentiating HHISs and LHSIs during the pre-induction period does not necessarily reflect expectation, but instead reflects this core component Our results add to the growing body of evidence relating EEG aperiodic activity in to inter-individual differences across diverse psychological research domains (Gerster et al., 2022; Pani et al., 2022). In particular, a new finding cognitive shifts in executive functioning during aging correlate with lower aperiodic exponent values (Finley et al., 2023). Such insights could be pivotal for understanding the relationship between aperiodic activity and hypnotic susceptibility, as cognitive control abilities are believed to play a significant role in modulating the hypnotic response (Egner & Raz, 2007; Faerman & Spiegel, 2021), while neuroimaging studies propose that this relationship encompasses prefrontal cortex activity (Cojan et al., 2015; Dienes & Hutton, 2013; Faerman et al., 2024). Nevertheless, the precise links between the aperiodic component of EEG recordings and executive functions are still underexplored, rendering these connections largely speculative at this point. Moreover, prevailing views propose that the aperiodic exponent indexes balance between excitatory and inhibitory activities of neural populations (Gao et al., 2017; Waschke et al., 2021). In this framework, a flatter aperiodic exponent corresponds to a greater excitation-inhibition ratio, which entails lower inhibition of cortical circuits and greater neuronal noise decoupled from oscillatory (synchronized). Prevailing views link such departure from inhibition and greater neuronal noise to poorer cognitive performance (Voytek & Knight, 2015). In this regard, at least one study lends support to the notion that baseline cortical excitability represents a neural characteristic of hypnotic susceptibility (Spina et al., 2020). Therefore, greater exponent values in HHSIs relative to LHSIs might reflect differences in executive control and cognitive control abilities associated with differences in neuronal excitability. These findings underscore the importance of pursuing further research that specifically examines the relationship between hypnotic susceptibility differences and the aperiodic component.

In conclusion, our research offers novel perspectives on the neural foundations of hypnotic susceptibility, and enriches the growing body of scientific literature highlighting the critical role of the aperiodic component in EEG signals for deciphering the nuances of neural differences between individuals (Bódizs et al., 2021; Cellier et al., 2021; Immink et al., 2021; Schaworonkow & Voytek, 2021; Waschke et al., 2017). Taken together, the outcome of our data-driven approach corroborates the idea that the hypnotic states, and our susceptibility to hypnotic suggestion, are mediated by several diverse electrophysiological components (Landry & Rainville, 2024). A surprising finding was the prominence of the exponent of the aperiodic EEG activity outside of hypnosis as the key neural feature. This observation supports the view that baseline brain activity offers an accurate representation of an individual’s susceptibility to hypnosis. Our findings pave the way for further research into the contribution of the aperiodic slope to the neural mechanisms that underlie hypnotic experiences, and more generally they demonstrate the feasibility of developing novel approaches to assess hypnotic susceptibility based on physiological recordings. Indeed, with the growing focus on the clinical use of hypnosis, it’s crucial to delve deeper into the variations in hypnotic response among individuals. This understanding is key to enhancing the effectiveness of clinical hypnosis within the framework of personalized medicine.

## Acknowledgments

M.L. acknowledges support from the Fonds de Recherche du Québec – Nature et Technologies, the Natural Science and Engineering Research Council of Canada. M.L., J.S. and A.R. acknowledge support from the Bial Foundation (Grant #280/16). J.d.C. acknowledges support from the Natural Science and Engineering Research Council of Canada. M.L., D.O. P.R. and K.J acknowledges support from the Fonds de Recherche du Québec – programme Audace (337355). K.J. is supported by funding from the Canada Research Chairs (950-232368) program and a Discovery Grant from the Natural Sciences and Engineering Research Council of Canada (2021-03426). We would also like to thank members of the HyMed lab and CoCo Lab.

## Contributions

Designed Research: All authors. Performed Research: M.L. and J.d.C. Funding acquisition: M.L., J.S., A.R., D.O., P.R. K.J. Contributed new analytic tools: M.L. and J.d.C. Analyzed Data: M.L. and J.d.C.; Wrote the Paper: All authors.

## Data availability

Data for the analyses supporting the current study is publicly available on the Open Science Framework repository:

## Code Availability

Code for the analyses supporting the current study is publicly available on the Open Science Framework repository:

## References

Baghdadi, G., & Nasrabadi, A. M. (2012). Comparison of different EEG features in estimation of hypnosis susceptibility level. Computers in biology and medicine, 42(5), 590–597.

Barnier, A. J., Cox, R. E., & McConkey, K. M. (2014). The Province of “Highs”: The High Hypnotizable Person in the Science of Hypnosis and in Psychological Science. Psychology of Consciousness: Theory, Research, and Practice, 1(2), 168–183.

Barnier, A. J., Terhune, D. B., Polito, V., & Woody, E. Z. (2022). A componential approach to individual differences in hypnotizability. *Psychology of Consciousness: Theory*, Research, and Practice, 9(2), 130.

Bódizs, R., Szalárdy, O., Horváth, C., Ujma, P. P., Gombos, F., Simor, P., Pótári, A., Zeising, M., Steiger, A., & Dresler, M. (2021). A set of composite, non-redundant EEG measures of NREM sleep based on the power law scaling of the Fourier spectrum. Scientific Reports, 11(1), 2041.

Cellier, D., Riddle, J., Petersen, I., & Hwang, K. (2021). The development of theta and alpha neural oscillations from ages 3 to 24 years. Developmental cognitive neuroscience, 50, 100969.

Cojan, Y., Piguet, C., & Vuilleumier, P. (2015). What makes your brain suggestible? Hypnotizability is associated with differential brain activity during attention outside hypnosis. NeuroImage, 117, 367–374.

Combrisson, E., & Jerbi, K. (2015). Exceeding chance level by chance: The caveat of theoretical chance levels in brain signal classification and statistical assessment of decoding accuracy. Journal of Neuroscience Methods, 250, 126–136.

Crawford, H. J., Clarke, S. W., & Kitner-Triolo, M. (1996, 1996/12/01/). Self-generated happy and sad emotions in low and highly hypnotizable persons during waking and hypnosis: laterality and regional EEG activity differences. International journal of psychophysiology, 24(3), 239–266.

De Pascalis, V. (1993). EEG spectral analysis during hypnotic induction, hypnotic dream and age regression. International journal of psychophysiology, 15(2), 153–166.

De Pascalis, V. (2023). EEG Oscillations and Neural Functional Connectivity Underpinning Hypnosis and Hypnotizability. Preprints.

De Pascalis, V. (2024a). Brain Functional Correlates of Resting Hypnosis and Hypnotizability: A Review. Brain Sciences, 14(2), 115.

De Pascalis, V. (2024b). EEG oscillatory activity concomitant with hypnosis and hypnotizability. In J. Linden, G. de Benedittis, L. Sugarman, & K. Varga (Eds.), The Routledge International Handbook of Clinical Hypnosis (pp. 244–255). Routledge.

De Pascalis, V., & Palumbo, G. (1986, 1986/02/01). EEG Alpha Asymmetry: Task Difficulty and Hypnotizability. Perceptual and motor skills, 62(1), 139–150.

De Pascalis, V., & Penna, P. M. (1990). 40-Hz Eeg Activity During Hypnotic Induction and Hypnotic Testing. International Journal of Clinical and Experimental Hypnosis, 38(2), 125–138.

Dienes, Z., & Hutton, S. (2013, 2//). Understanding hypnosis metacognitively: rTMS applied to left DLPFC increases hypnotic suggestibility. Cortex, 49(2), 386–392.

Dietterich, T. G. (1998). Approximate statistical tests for comparing supervised classification learning algorithms. Neural computation, 10(7), 1895–1923.

Donoghue, T., Haller, M., Peterson, E. J., Varma, P., Sebastian, P., Gao, R., Noto, T., Lara, A. H., Wallis, J. D., Knight, R. T., Shestyuk, A., & Voytek, B. (2020). Parameterizing neural power spectra into periodic and aperiodic components. Nature neuroscience, 23(12), 1655–1665.

Donoghue, T., Schaworonkow, N., & Voytek, B. (2022). Methodological considerations for studying neural oscillations. European Journal of Neuroscience, 55(11-12), 3502–3527.

Egner, T., & Raz, A. (2007). Cognitive Control Processes and Hypnosis. In G. A. Jamieson (Ed.), Hypnosis and Conscious States: The cognitive neuroscience perspective (pp. 29–50). Oxford University Press.

Faerman, A., Bishop, J. H., Stimpson, K. H., Phillips, A., Gülser, M., Amin, H., Nejad, R., DeSouza, D. D., Geoly, A. D., Kallioniemi, E., Jo, B., Williams, N. R., & Spiegel, D. (2024, 2024/01/01). Stanford Hypnosis Integrated with Functional Connectivity-targeted Transcranial Stimulation (SHIFT): a preregistered randomized controlled trial. Nature Mental Health, 2(1), 96–103.

Faerman, A., & Spiegel, D. (2021). Shared cognitive mechanisms of hypnotizability with executive functioning and information salience. Scientific Reports, 11(1), 5704.

Finley, A. J., Angus, D. J., Knight, E., van Reekum, C. M., Lachman, M. E., Davidson, R. J., & Schaefer, S. M. (2023). Resting EEG Periodic and Aperiodic Components Predict Cognitive Decline Over 10 Years. bioRxiv, 2023-2007.

Gandhi, B., & Oakley, D. A. (2005). Does ‘hypnosis’ by any other name smell as sweet? The efficacy of ‘hypnotic’inductions depends on the label ‘hypnosis’. Consciousness and cognition, 14(2), 304–315.

Gao, R., Peterson, E. J., & Voytek, B. (2017). Inferring synaptic excitation/inhibition balance from field potentials. NeuroImage, 158, 70–78.

Gerster, M., Waterstraat, G., Litvak, V., Lehnertz, K., Schnitzler, A., Florin, E., Curio, G., & Nikulin, V. (2022). Separating neural oscillations from aperiodic 1/f activity: challenges and recommendations. Neuroinformatics, 1-22.

Graffin, N. F., Ray, W. J., & Lundy, R. (1995). EEG concomitants of hypnosis and hypnotic susceptibility. Journal of Abnormal Psychology, 104(1), 123.

Gyurkovics, M., Clements, G. M., Low, K. A., Fabiani, M., & Gratton, G. (2022). Stimulus-induced changes in 1/f-like background activity in EEG. Journal of Neuroscience, 42(37), 7144–7151.

Haufe, S., Meinecke, F., Görgen, K., Dähne, S., Haynes, J.-D., Blankertz, B., & Bießmann, F. (2014). On the interpretation of weight vectors of linear models in multivariate neuroimaging. NeuroImage, 87, 96–110.

Haynes, J.-D., & Rees, G. (2006, 2006/07/01). Decoding mental states from brain activity in humans. Nature Reviews Neuroscience, 7(7), 523–534.

He, B. J. (2014). Scale-free brain activity: past, present, and future. Trends in cognitive sciences, 18(9), 480–487.

Hoeft, F., Gabrieli, J. D., Whitfield-Gabrieli, S., Haas, B. W., Bammer, R., Menon, V., & Spiegel, D. (2012). Functional brain basis of hypnotizability. Archives of General Psychiatry, 69(10), 1064–1072.

Huber, A., Lui, F., Duzzi, D., Pagnoni, G., & Porro, C. A. (2014). Structural and Functional Cerebral Correlates of Hypnotic Suggestibility. PloS one, 9(3), e93187.

Immink, M. A., Cross, Z. R., Chatburn, A., Baumeister, J., Schlesewsky, M., & Bornkessel-Schlesewsky, I. (2021). Resting-state aperiodic neural dynamics predict individual differences in visuomotor performance and learning. Human Movement Science, 78, 102829.

Jamieson, G. A. (2007). Hypnosis and conscious states: The cognitive neuroscience perspective. Oxford University Press.

Jamieson, G. A., & Burgess, A. P. (2014). Hypnotic induction is followed by state-like changes in the organization of EEG functional connectivity in the theta and beta frequency bands in high-hypnotically susceptible individuals. Frontiers in human neuroscience, 8, 528.

Jensen, M. P., Adachi, T., & Hakimian, S. (2015). Brain Oscillations, Hypnosis, and Hypnotizability. American Journal of Clinical Hypnosis, 57(3), 230–253.

Kumar, V. K., & Pekala, R. J. (1989). Variations in the phenomenological experience as a function of hypnosis and hypnotic susceptibility: A replication. British Journal of Experimental & Clinical Hypnosis.

Landry, M., Lifshitz, M., & Raz, A. (2017). Brain correlates of hypnosis: A systematic review and meta-analytic exploration. Neuroscience & Biobehavioral Reviews, 81, 75–98.

Landry, M., & Rainville, P. (2024). Beyond the neural signature of hypnosis: Neuroimaging studies support a multifaceted view of hypnotic phenomena. In J. Linden, L. Sugerman, G. de Benedittis, & K. Varga (Eds.), Routledge International Handbook of Clinical Hypnosis. Taylor & Francis.

Landry, M., & Raz, A. (2015). Hypnosis and imaging of the living brain. American Journal of Clinical Hypnosis, 57(3), 285–313.

Landry, M., & Raz, A. (2016). Heightened states of attention: From mental performance to altered states of consciousness and contemplative practices. Intellectica, 66, 139–159.

Lardone, A., Liparoti, M., Sorrentino, P., Minino, R., Polverino, A., Lopez, E. T., Bonavita, S., Lucidi, F., Sorrentino, G., & Mandolesi, L. (2022). Topological changes of brain network during mindfulness meditation: an exploratory source level magnetoencephalographic study. AIMS Neuroscience, 9(2), 250–263.

Latora, V., & Marchiori, M. (2001). Efficient Behavior of Small-World Networks. Physical Review Letters, 87(19), 198701.

Laurence, J.-R., Beaulieu-Prévost, D., & Du Chéné, T. (2008). Measuring and understanding individual differences in hypnotizability. In A. J. Barnier & M. R. Nash (Eds.), The Oxford Handbook of Hypnosis: Theory, Research, and Practice. Oxford University Press.

Lee, J.-S., Spiegel, D., Kim, S.-B., Lee, J.-H., Kim, S.-I., Yang, B.-H., Choi, J.-H., Kho, Y.-C., & Nam, J.-H. (2007). Fractal analysis of EEG in hypnosis and its relationship with hypnotizability. Intl. Journal of Clinical and Experimental Hypnosis, 55(1), 14–31.

Lifshitz, M., Aubert Bonn, N., Fischer, A., Kashem, I. F., & Raz, A. (2013, Feb). Using suggestion to modulate automatic processes: from Stroop to McGurk and beyond. Cortex, 49(2), 463–473.

Lifshitz, M., van Elk, M., & Luhrmann, T. M. (2019). Absorption and spiritual experience: A review of evidence and potential mechanisms. Consciousness and cognition, 73, 102760.

Lopes da Silva, F. (2013, 2013/12/04/). EEG and MEG: Relevance to Neuroscience. Neuron, 80(5), 1112–1128.

Maschke, C., Duclos, C., Owen, A. M., Jerbi, K., & Blain-Moraes, S. (2023). Aperiodic brain activity and response to anesthesia vary in disorders of consciousness. NeuroImage, 275, 120154.

McNemar, Q. (1947). Note on the sampling error of the difference between correlated proportions or percentages. Psychometrika, 12(2), 153–157.

Newman, M. E. (2006). Modularity and community structure in networks. Proceedings of the national academy of sciences, 103(23), 8577–8582.

Onnela, J.-P., Saramäki, J., Kertész, J., & Kaski, K. (2005). Intensity and coherence of motifs in weighted complex networks. Physical Review E, 71(6), 065103.

Panda, R., Vanhaudenhuyse, A., Piarulli, A., Annen, J., Demertzi, A., Alnagger, N., Chennu, S., Laureys, S., Faymonville, M.-E., & Gosseries, O. (2023). Altered Brain Connectivity and Network Topological Organization in a Non-ordinary State of Consciousness Induced by Hypnosis. Journal of Cognitive Neuroscience, 35(9), 1394–1409.

Pani, S. M., Saba, L., & Fraschini, M. (2022). Clinical applications of EEG power spectra aperiodic component analysis: A mini-review. Clinical Neurophysiology.

Pei, L., Zhou, X., Leung, F. K. S., & Ouyang, G. (2023). Differential associations between scale-free neural dynamics and different levels of cognitive ability. Psychophysiology, 60(6), e14259.

Pekala, R. J. (2015). Hypnosis as a “state of consciousness”: How quantifying the mind can help us better understand hypnosis. American Journal of Clinical Hypnosis, 57(4), 402–424.

Rainville, P., & Price, D. D. (2003). Hypnosis phenomenology and the neurobiology of consciousness. International Journal of Clinical and Experimental Hypnosis, 51(2), 105–129.

Raz, A., & Lifshitz, M. (2016). Hypnosis and meditation: Towards an integrative science of conscious planes. Oxford University Press.

Rho, G., Callara, A. L., Petri, G., Nardelli, M., Scilingo, E. P., Greco, A., & Pascalis, V. D. (2021). Linear and Nonlinear Quantitative EEG Analysis during Neutral Hypnosis following an Opened/Closed Eye Paradigm. Symmetry, 13(8).

Rubinov, M., & Sporns, O. (2010). Complex network measures of brain connectivity: uses and interpretations. NeuroImage, 52(3), 1059–1069.

Rubinov, M., & Sporns, O. (2011). Weight-conserving characterization of complex functional brain networks. NeuroImage, 56(4), 2068–2079.

Schaworonkow, N., & Voytek, B. (2021). Longitudinal changes in aperiodic and periodic activity in electrophysiological recordings in the first seven months of life. Developmental cognitive neuroscience, 47, 100895.

Shor, R. E., & Orne, E. C. (1962). Harvard group scale of hypnotic susceptibility, Form: A. Consulting psychologists press.

Spiegel, D., Cutcomb, S., Ren, C., & Pribram, K. (1985). Hypnotic hallucination alters evoked potentials. Journal of Abnormal Psychology, 94(3), 249.

Spina, V., Chisari, C., & Santarcangelo, E. L. (2020). High motor cortex excitability in highly hypnotizable individuals: a favourable factor for neuroplasticity? Neuroscience, 430, 125–130.

Sugimoto, K., Kurashiki, H., Xu, Y., Takemi, M., & Amano, K. (2024). Electroencephalographic Biomarkers of Relaxation: A Systematic Review and Meta-analysis. bioRxiv, 2024-2003.

Tadel, F., Baillet, S., Mosher, J. C., Pantazis, D., & Leahy, R. M. (2011). Brainstorm: a user-friendly application for MEG/EEG analysis. Computational intelligence and neuroscience, 2011, 8.

Taghilou, H., Rezaei, M., Nazari, M. A., & Valizadeh, A. (In press). EEG Oscillations during Prehypnosis and Hypnosis in Subjects with High and Low Dissociative Experiences. Basic and clinical neuroscience.

Terhune, D. B., & Cardeña, E. (2016). Nuances and Uncertainties Regarding Hypnotic Inductions: Toward a Theoretically Informed Praxis. American Journal of Clinical Hypnosis, 59(2), 155–174.

Terhune, D. B., Cleeremans, A., Raz, A., & Lynn, S. J. (2017). Hypnosis and top-down regulation of consciousness. Neuroscience & Biobehavioral Reviews.

Thölke, P., Mantilla-Ramos, Y.-J., Abdelhedi, H., Maschke, C., Dehgan, A., Harel, Y., Kemtur, A., Berrada, L. M., Sahraoui, M., & Young, T. (2022). Class imbalance should not throw you off balance: Choosing the right classifiers and performance metrics for brain decoding with imbalanced data. bioRxiv, 2022-2007.

Timmermann, C., Bauer, P. R., Gosseries, O., Vanhaudenhuyse, A., Vollenweider, F., Laureys, S., Singer, T., Antonova, E., & Lutz, A. (2023). A neurophenomenological approach to non-ordinary states of consciousness: hypnosis, meditation, and psychedelics. Trends in cognitive sciences, 27(2), 139–159.

van Lutterveld, R., van Dellen, E., Pal, P., Yang, H., Stam, C. J., & Brewer, J. (2017). Meditation is associated with increased brain network integration. NeuroImage, 158, 18–25.

Vanhaudenhuyse, A., Laureys, S., & Faymonville, M. E. (2014). Neurophysiology of hypnosis. Neurophysiologie Clinique/Clinical Neurophysiology, 44(4), 343–353.

Vinck, M., Oostenveld, R., van Wingerden, M., Battaglia, F., & Pennartz, C. M. A. (2011). An improved index of phase-synchronization for electrophysiological data in the presence of volume-conduction, noise and sample-size bias. NeuroImage, 55(4), 1548–1565.

Voytek, B., & Knight, R. T. (2015). Dynamic network communication as a unifying neural basis for cognition, development, aging, and disease. Biological psychiatry, 77(12), 1089–1097.

Waschke, L., Donoghue, T., Fiedler, L., Smith, S., Garrett, D. D., Voytek, B., & Obleser, J. (2021). Modality-specific tracking of attention and sensory statistics in the human electrophysiological spectral exponent. Elife, 10, e70068.

Waschke, L., Wöstmann, M., & Obleser, J. (2017). States and traits of neural irregularity in the age-varying human brain. Scientific Reports, 7(1), 17381.

Weitzenhoffer, A. M. (1980). Hypnotic susceptibility revisited. American Journal of Clinical Hypnosis, 22(3), 130–146.

Williams, J. D., & Gruzelier, J. H. (2001, 2001/07/01). Differentiation of hypnosis and relaxation by analysis of narrow band theta and alpha frequencies. International Journal of Clinical and Experimental Hypnosis, 49(3), 185–206.

Wilson, L. E., da Silva Castanheira, J., & Baillet, S. (2022). Time-resolved parameterization of aperiodic and periodic brain activity. Elife, 11, e77348.

Yapko, M. D. (2015). Essentials of hypnosis (2nd ed.). Routledge.

