## supplementary material for "Aperiodic activity as a central neural feature of hypnotic susceptibility outside of hypnosis"

We present values derived from spectral features and graph theoretical clustering coefficients, which are based on wPLI connectivity patterns. Following our comprehensive multivariate pattern analysis, three EEG channels were strategically selected for each neural feature where we were able to classify hypnotic susceptibility better than chance level. We detail the values for these channels during both the pre-induction and post-induction periods, as well as the differential values between these two states. Values are presented separately for LHSIs and HHSIs. Consequently, this illustration facilitates the tracking of variations across different conditions, thereby providing a deeper insight into the implications and effectiveness of our classification procedure.

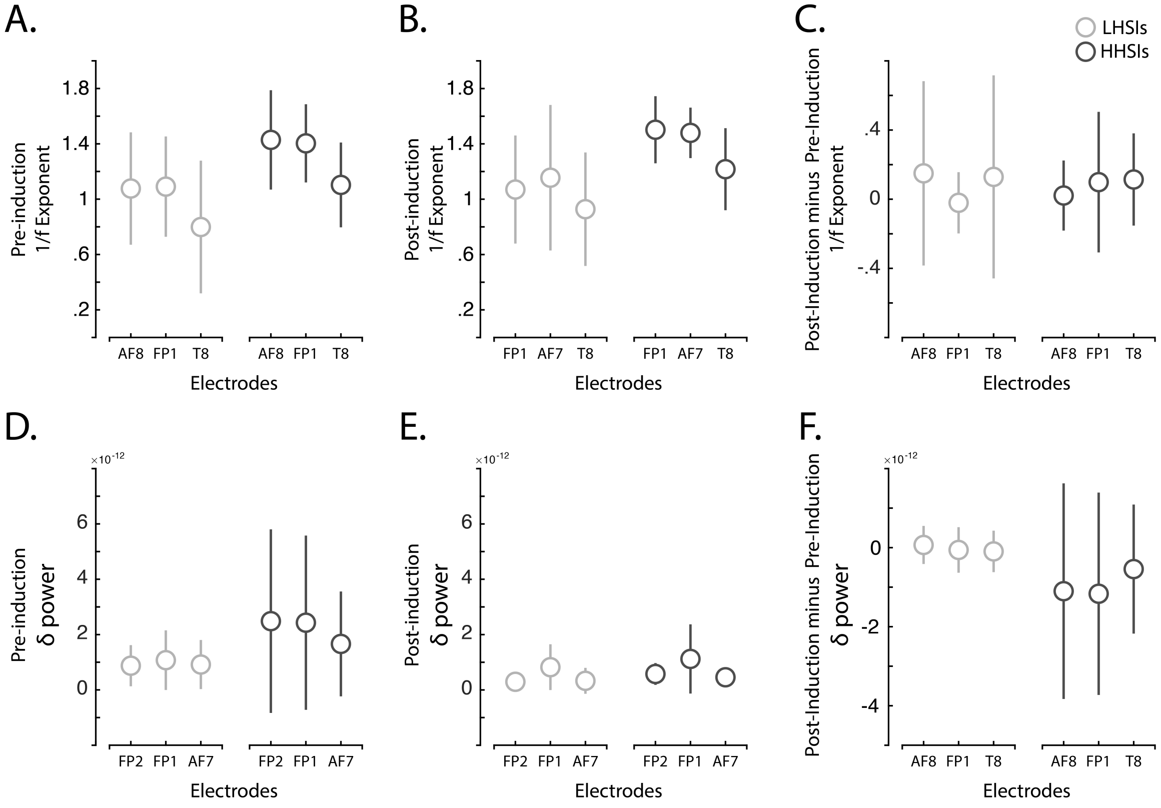


**Supplementary Figure 1.** Estimations of exponent slope values for channels AF8, FP1, and T8 across Low Hypnotic Susceptibility Individuals (LHSIs) and High Hypnotic Susceptibility Individuals (HHSIs). Panel A illustrates these estimates for the pre-induction period, Panel B for the post-induction period, and Panel C details the variance between these two conditions. Additionally, the figure presents spectral power data for broadband delta frequencies in channels FP2, FP1, and AF7. This data is also segmented across LHSIs and HHSIs, with Panel D showing the pre-induction period, Panel E the post-induction period, and Panel F highlighting the differences between these conditions.

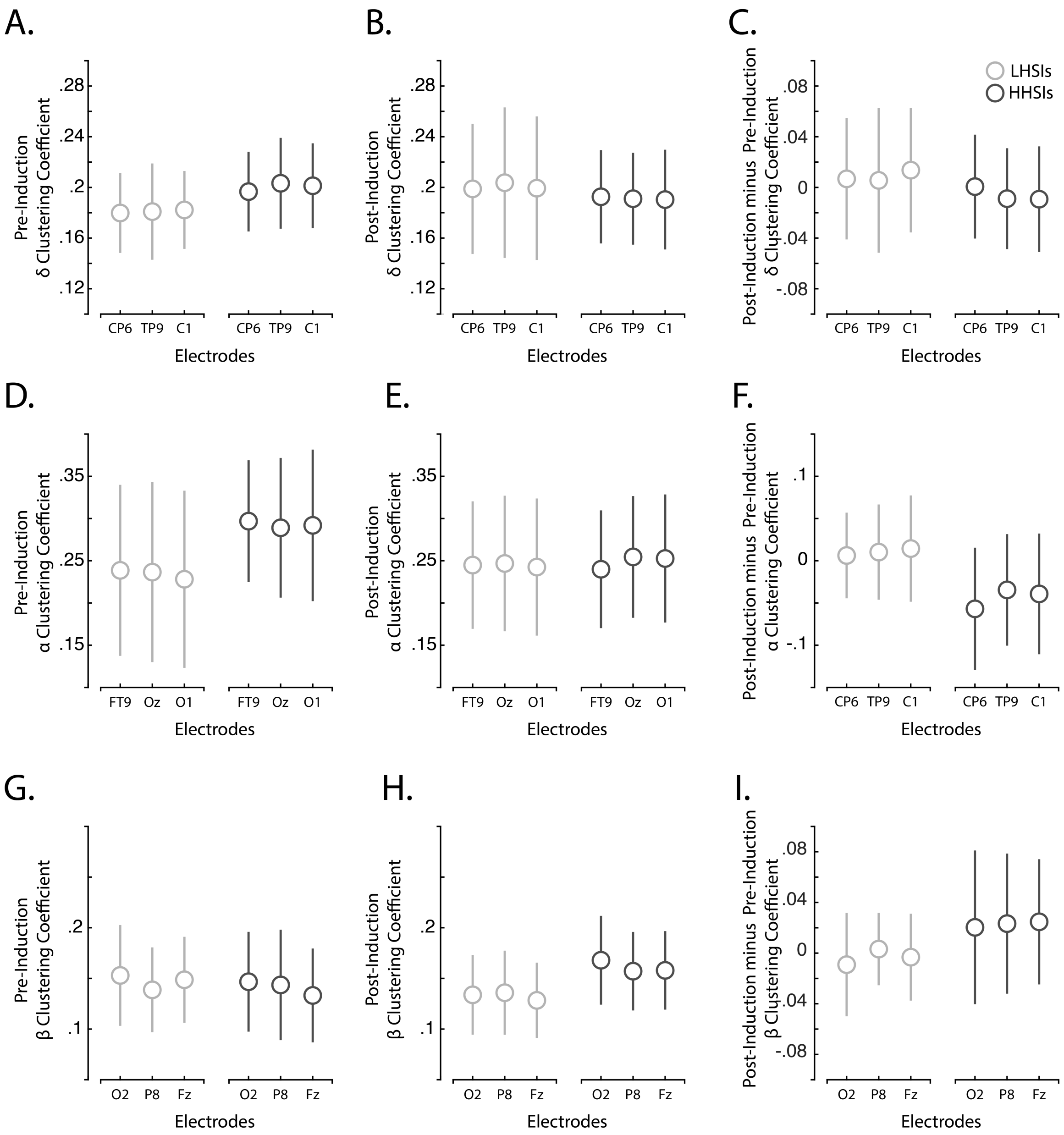


**Supplementary Figure 2.** Estimations of clustering coefficients across delta (panels A, B, C; channels CP6, TP9, and C1), alpha (panels D, E, F; channels FT9, Oz, and O1), and beta (panels G, H, I; channels O2, P8, and Fz) frequencies across Low Hypnotic Susceptibility Individuals (LHSIs) and High Hypnotic Susceptibility Individuals (HHSIs). Panels A, D, and G correspond to the pre-induction period, panels B, E and H the post-induction period, and Panel C, F and I indicate the differences between these conditions.

We used model comparison to identify the top feature for predicting hypnotic susceptibility. We used McNemar's chi-square statistics to compare the performance of models (**Supplementary figure 3**).

B.

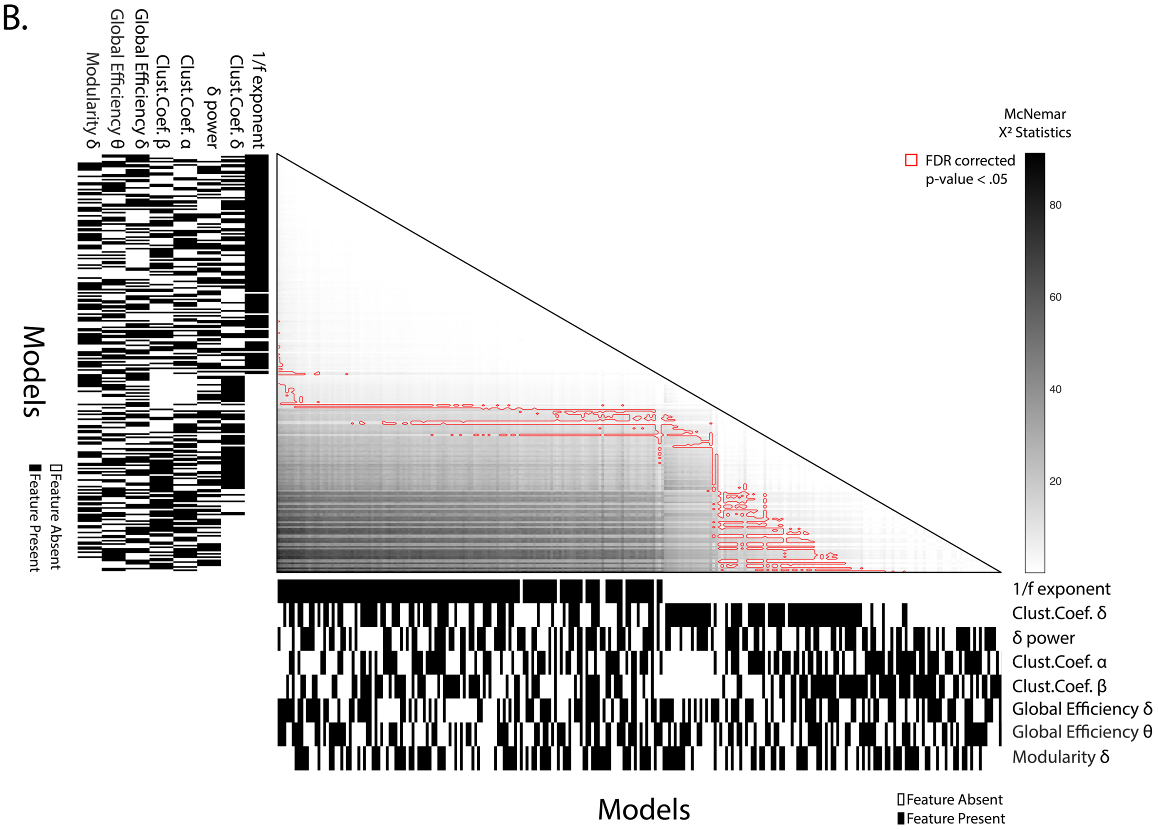


**Supplementary Figure 3.** Panel A shows the corresponding McNemar's Chi-square statistics where we compared all models in a pairwise manner. Red contours indicate statistically significant comparisons after FDR corrections for multiple comparisons.

Next, here is the script we used for the hypnotic induction across all participants.

**Hypnotic Induction Script**

Alright, let's begin.

Right now, there is nothing to do at all.
Your mind might start to wander…
You might feel the chair below you, or the EEG cap on your head…
and as you notice these sensations...
you might find yourself noticing the sound of my voice…
and as you notice the sound of my voice...
or notice your sensations…
You may also begin to become aware of how easy it would be to close your eyes right now…
And as you begin to wonder what it would feel like to close your eyes now…
As you continue to notice the sound of my voice…
your eyelids might feel heavy…
Or perhaps they might just feel tired…
I don't know to what extent your eyelids might feel heavy or tired…

However they feel, these sensations might make you want to just
close your eyes….
Perhaps you can imagine heavy eyes, tired eyes…
and as you notice this...
it may feel nice to just let them rest.
And as you let your eyes rest now, you might imagine that you are already developing the capacity to enter a very deep state, now…
And as my voice continues, you may also become aware of your breathing…
In-breath, and out-breath
Perhaps you can imagine that with each out-breath, you go deeper and deeper ... more relaxed, more deep
And with each out-breath you may notice yourself going deeper now (time with outbreaths)
That's right
Perhaps you feel relaxed right now...
perhaps you feel heavy…
Perhaps you feel in a deep, deep state...
Anyway you feel right now is exactly right
And as you are aware of how you feel,
Deep, tired, heavy…
as you become aware of how you feel..
you might notice you also grow more aware of your capacity to enter a very deep state…
Somehow this might remind you of other capacities you have to enter different mental states….
perhaps this might remind you of a specific mental state….
A time when you couldn't read…
Maybe this might be when looking at words in another language…
Or perhaps this might remind you of when you were a child, before you learned to read....
And as you remember this… you might notice that written words begin to change in your minds eye…
Perhaps they simply look 'off', strange…
Or perhaps they appear to be written in a foreign language…
However written words change for you now is perfectly fine...
As my voice continues…
You might notice your relationship to written words continues to change…
so that words might become completely illegible....
you might imagine that if you saw a written word in front of you…
You might not be able to read it…
and perhaps as you imagine being completely unable to read words in front of you…
You might notice that other abilities are preserved….
Numbers in numerical form are completely legible…
And as you continue to notice how strange and illegible written words appear,
You might grow increasingly aware of how clear and legible numbers are.

And you might notice the contrast between numbers that are spelled out…
that might be blurry, illegible, confusing, nonsensical...
And numbers that appear in numerical form…
clear, crystal clear...
And you can just allow this process to naturally happen…
you can rest assured that your body and mind
Know how to do that for you …
And will continue to do so now…
And just allow this experience to happen…
you can simply allow the experience of being unable to read words arise …
However it may arise…
That’s right…
As my voice continues…
You may notice how deep your mental state is...
and as you notice this...
you might notice your capacity to change how you read deepen…
That’s right.

So now we will begin the first task.
